# Claudin-1 Inhibitor PDS-0330 Ameliorates Diabetic Kidney Disease by Suppressing Src/Akt/mTOR Signaling Pathway in Podocytes

**DOI:** 10.64898/2025.12.24.696454

**Authors:** Kazuhiko Fukushima, Kenji Tsuji, Hiroyuki Nakanoh, Naruhiko Uchida, Shinji Kitamura, Jun Wada

**Affiliations:** Department of Nephrology, Rheumatology, Endocrinology and Metabolism, Okayama University Graduate School of Medicine, Dentistry and Pharmaceutical Sciences, 2-5-1 Shikata-cho, Okayama, 700-8558 Japan

**Keywords:** claudin-1, diabetic kidney disease, podocyte

## Abstract

**Introduction:** Recent studies have reported that claudin-1, which is reactively expressed in malignant tumors and inflammatory diseases, plays an important role in disease progression. In diabetic kidney disease (DKD), upregulated expression of claudin-1 in podocytes has also been shown to contribute to disease progression. However, it remains unclear whether claudin-1 itself could be a therapeutic target in DKD. In this study, we investigated whether the claudin-1 inhibitor PDS-0330 could ameliorate DKD.

**Methods:** Five-week-old male C57BL/6J mice were fed either a high-fat diet (HFD) or a normal diet (ND). At 15 weeks of age, PDS-0330 (Inh) or vehicle as control (Ctrl) was administered orally once per week for 4 weeks. At the end of the treatment period, blood, urine, and kidney samples were collected. In *in vitro* study, immortalized podocytes were treated with Inh or Ctrl and cultured for 48 hours in either normal glucose medium (5.5 mM; NG) or high glucose medium (25 mM; HG), after which proteins were extracted for analysis.

**Results:** Mice fed an HFD showed significant increases in body weight and serum glucose levels compared with ND mice, and these parameters were not significantly affected by Inh treatment. HFD mice treated with Ctrl (HFD-Ctrl) exhibited significantly higher urinary albumin excretion than ND mice treated with Ctrl (ND-Ctrl), whereas the Inh-treated HFD mice (HFD-Inh) showed a significant reduction. Transmission electron microscopy revealed foot process effacement in HFD-Ctrl podocytes, which was markedly improved in HFD-Inh mice. Immunofluorescence staining demonstrated claudin-1 expression in podocytes of HFD mice. Furthermore, phospho-mTOR staining in podocytes was significantly increased in HFD-Ctrl compared with ND-Ctrl and was significantly attenuated in HFD-Inh. Western blot analysis of kidney samples revealed activation of the Src/Akt/mTOR signaling pathway in HFD-Ctrl mice, which was significantly reduced in HFD-Inh mice. In *in vitro* study, Ctrl-treated podocytes cultured in HG (HG-Ctrl) showed significant activation of Src/Akt/mTOR signaling compared with those in NG (NG-Ctrl). This activation was significantly suppressed in Inh-treated podocytes (HG-Inh).

**Conclusion:** These findings suggest that podocyte claudin-1 could be a potential therapeutic target in DKD.

## Introduction

Diabetic kidney disease (DKD) is one of the most important complications of diabetes mellitus, as it contributes to cardiovascular disease and is one of the leading causes of end-stage renal disease. Various therapeutic agents for glycemic control have been developed over the past several decades, however, far fewer drugs targeting DKD have been developed. One reason for this lag is that the pathogenic mechanisms underlying the progression of DKD have not been fully elucidated. In the early stages of DKD, the condition can improve reversibly with appropriate control of blood glucose and blood pressure, but in advanced stages, it deteriorates irreversibly[1]. Thus, researchers worldwide are working to clarify the mechanisms driving DKD progression and to develop new therapeutic strategies.

Increasing attention has been focused on the involvement of claudin-1 in the progression of DKD. Claudin-1 is well known as a major component of tight junctions and a molecule essential for intercellular adhesion[2]. However, under pathological conditions such as inflammatory diseases and cancers, claudin-1 is reactively expressed in a form referred to as non-junctional claudin-1, which does not form tight junctions but instead has been reported to amplify signaling pathways that promote cell proliferation and migration[3]. In fact, there have been reports demonstrating that anti–claudin-1 antibody therapeutics exert therapeutic effects on liver cirrhosis, pulmonary fibrosis, and hepatocellular carcinoma[4] [5]. In the kidney field as well, there is a report showing that anti–claudin-1 antibody therapy ameliorates interstitial fibrosis in the unilateral ureteral obstruction (UUO) model, although the study did not specify which renal cell types – such as tubular epithelial cells or interstitial fibroblasts – were directly targeted by the antibody[4]. A clinical study evaluating the therapeutic efficacy of the anti–claudin-1 antibody in crescentic glomerulonephritis is currently underway (RENAL-F02, NCT06047171), indicating that claudin-1 targeting is already being explored in the clinical setting. In the study, the anti–claudin-1 antibody is intended to target parietal epithelial cells of Bowman’s capsule, which constitutively express claudin-1 and represent the principal cellular component of crescent formation. In DKD, reactive expression of claudin-1 in podocytes is known to play a critical role in disease progression; specifically, decreased SIRT1 expression in proximal tubular cells promotes claudin-1 expression in podocytes, thereby increasing urinary protein excretion[6] [7]. These reports suggest that claudin-1 expressed in podocytes interferes with the renoprotective effects of the SIRT1 signaling pathway. However, it remains unclear whether targeting claudin-1 that is reactively induced in podocytes could be a therapeutic strategy, given that the biological behavior of claudin-1 in podocytes may differ from that of claudin-1 constitutively expressed in parietal epithelial cells.

More recently, PDS-0330, an inhibitor of claudin-1–mediated Src activation, has been developed[8]. PDS-0330 was originally created as an anticancer agent, suppressing claudin-1–induced Src activation in malignant tumor cells, a process that triggers tumor cell proliferation and metastasis. Src signaling has also been reported to play an important role in the progression of DKD, and indeed, several studies have shown that Src inhibitors can attenuate DKD[9]. Based on this background, we investigated whether PDS-0330 exerts therapeutic effects in DKD.

## Results

### Claudin-1 inhibitor PDS-0330 reduces urinary albumin excretion in a diabetic mouse model

As described in the Materials and Methods, five-week-old male C57BL/6J mice were fed either a normal diet (ND) or a high-fat diet (HFD) for 10 weeks, followed by administration of PDS-0330 (5 mg/kg) or control vehicle for 4 weeks (Fig. 1A). We determined the dose of PDS-0330 based on the previous report and concluded that a dose of 5 mg/kg in mice effectively inhibits claudin-1–dependent Src activation without causing adverse effects such as weight loss[8]. As shown in Fig. 1B, HFD mice exhibited a significant increase in body weight compared with ND mice, whereas neither PDS-0330 nor vehicle treatment resulted in significant changes in body weight. Serum analysis demonstrated significantly elevated serum glucose levels in HFD mice compared with ND mice (Fig. 2A). In both ND and HFD mice, PDS-0330 administration did not induce any significant changes in serum glucose levels, urea nitrogen levels or creatinine levels (Fig. 2A-C). Urinary albumin excretion in HFD mice was significantly higher than that in ND mice (Fig. 2D). Interestingly, HFD mice treated with PDS-0330 showed a significant reduction in urinary albumin (Fig. 2D). Unexpectedly, PDS-0330 also significantly reduced urinary albumin excretion in ND mice (Fig. 2D) although urinary albumin levels in ND mice were consistent with those reported previously[10, 11]. These findings suggest that PDS-0330 reduces albuminuria in both HFD and ND mice via mechanisms independent of blood glucose or body weight control.

**Figure 1.**
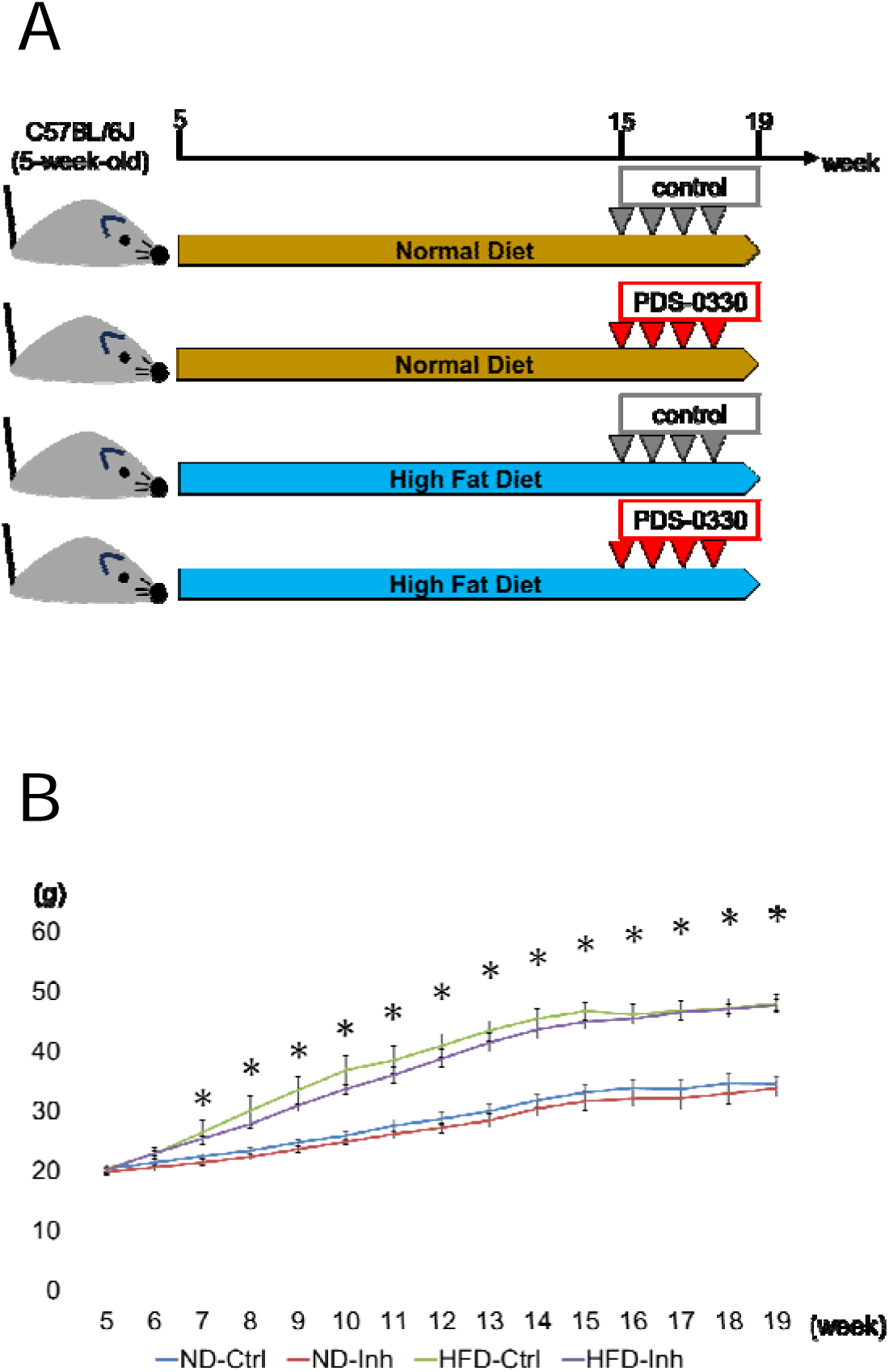
Schedule of diet feeding and claudin-1 inhibitor administration in mice, and corresponding body weight changes. Normal diet (ND)-fed or high-fat diet (HFD)-fed mice were orally administered either the claudin-1 inhibitor PDS-0330 (Inh) or a control solvent (Ctrl) once weekly from 15 weeks of age for 4 weeks. (A) Timeline of mouse feeding and drug administration. (B) Body weight changes during diet feeding and Inh administration. Results are presented as the mean ± standard deviation. Differences were evaluated by two-way ANOVA followed by the Tukey–Kramer test; * p < 0.05 for ND-Ctrl vs. HFD-Ctrl.

**Figure 2.**
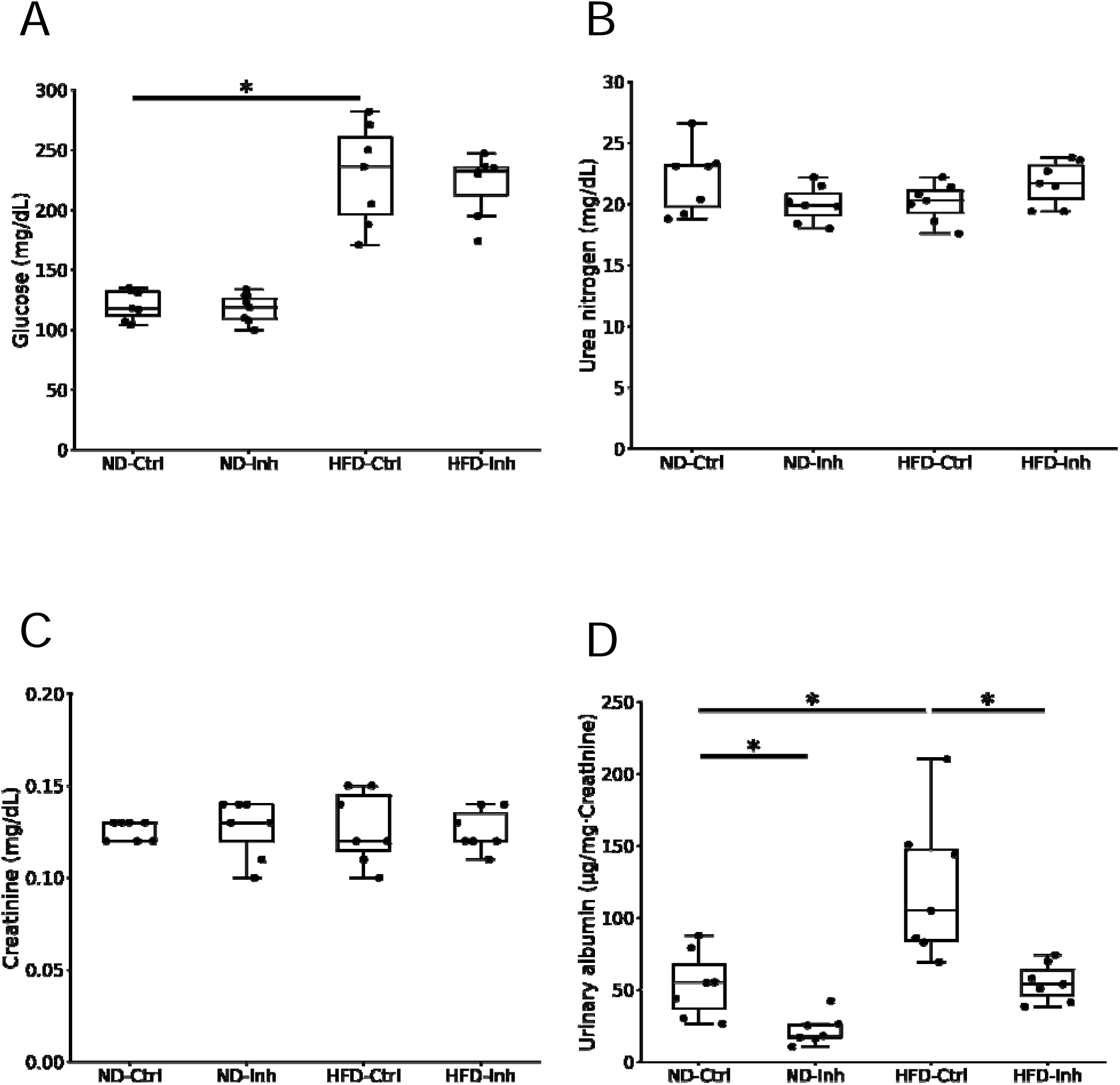
Biochemical and urinary parameters in mice fed a normal or high-fat diet and treated with or without claudin-1 inhibitor. Serum samples were collected at the time of sacrifice, and urine samples were obtained from a 24-hour urine collection performed on the day prior to sacrifice. (A) Serum glucose. (B) Serum urea nitrogen. (C) Serum creatinine. (D) Urinary albumin. Differences were evaluated by two-way ANOVA followed by the Tukey–Kramer test; * p < 0.05. ND, normal diet; HFD, high-fat diet; Ctrl, control solvent; Inh, claudin-1 inhibitor PDS-0330.

### Claudin-1 inhibitor PDS-0330 protects podocytes in a diabetic mouse model

We next examined the effects of PDS-0330 on renal histopathology. Consistent with previous reports[12, 13], PAS staining revealed significant glomerular enlargement and expansion of the mesangial area in HFD mice compared with ND mice (Fig. 3A). Transmission electron microscopy (TEM) also demonstrated mesangial matrix expansion and increased mesangial expansion in HFD mice (Fig. 3B). In addition, glomerular capillary loops showed narrowing of the slit diaphragm and foot process effacement in HFD mice. Interestingly, HFD mice treated with PDS-0330 displayed significant improvement in these podocyte-associated abnormalities (Fig. 3C), findings that paralleled the reduction in urinary albumin excretion (Fig. 2D). These observations suggest that PDS-0330 exerts protective effects on podocytes in HFD mice.

**Figure 3.**
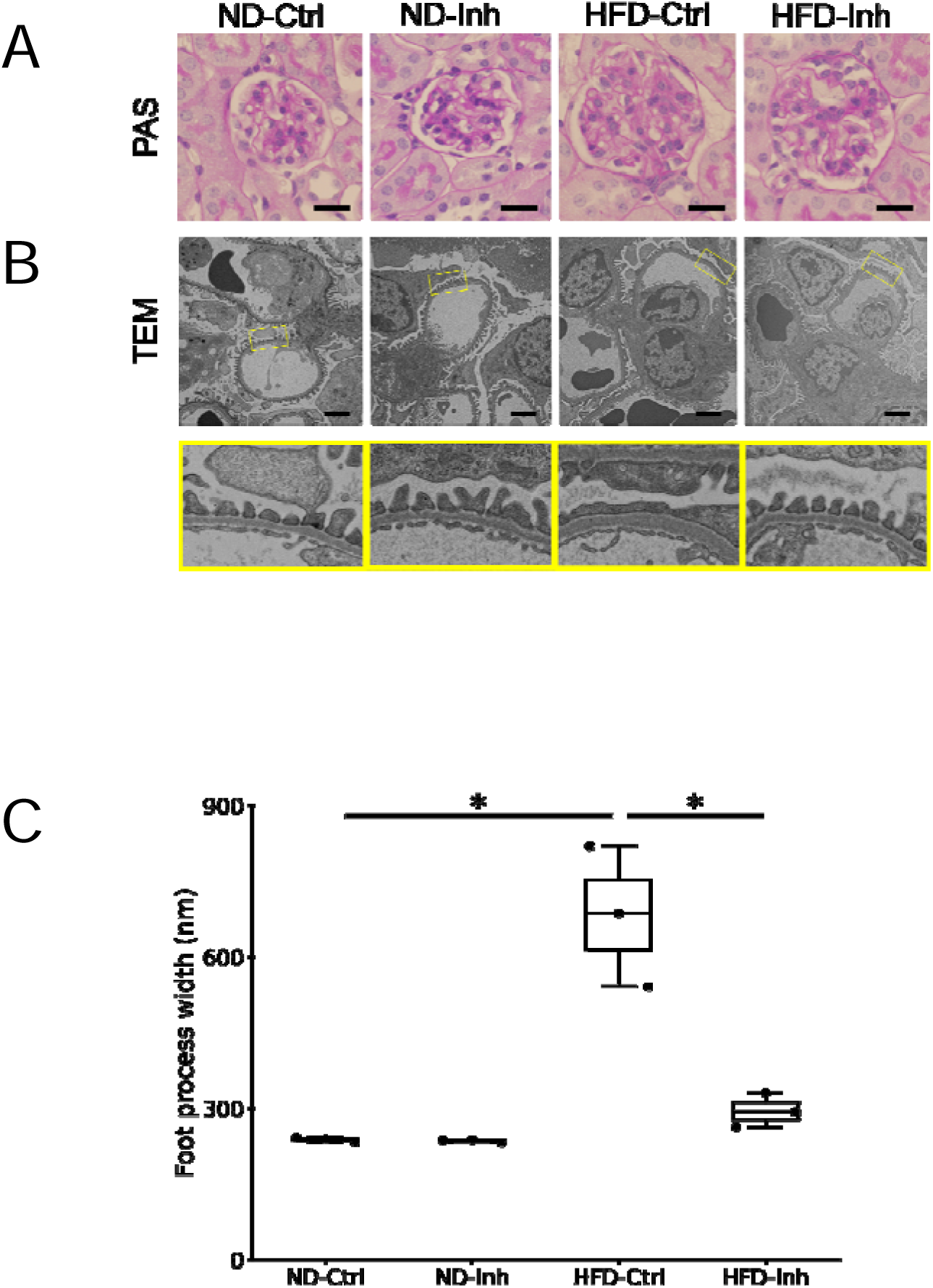
Histological and ultrastructural analysis of glomeruli in mice with or without claudin-1 inhibitor Treatment. (A) Periodic Acid–Schiff staining. Scale bars = 20 μm. (B) Transmission electron microscopy (TEM). Scale bars = 2.0 µm. (C) Podocyte foot process width measurement based on transmission electron microscopy. Differences were evaluated by two-way ANOVA followed by the Tukey–Kramer test; * p < 0.05. ND, normal diet; HFD, high-fat diet; Ctrl, control solvent; Inh, claudin-1 inhibitor PDS-0330.

### Claudin-1 inhibitor PDS-0330 attenuates Src/Akt/mTOR signaling activation in podocytes of diabetic mouse models

We next investigated the mechanisms underlying the protective effects on podocytes in DKD by PDS-0330. PDS-0330 has been reported to inhibit claudin-1–mediated Src signaling activity in colorectal cancer cells[8]. Meanwhile, in DKD, claudin-1 is reactively expressed in podocytes and contributes to disease progression[6]. Previous studies have also demonstrated Src activation in DKD and reported that pharmacological Src inhibition ameliorates the disease[9]. Src is known to influence various downstream signaling pathways, including the Akt/mTOR pathway, which plays a major role in the progression of DKD[14–16]. Based on these backgrounds, we evaluated Src/Akt/mTOR signaling in mouse renal cortical samples using immunofluorescence staining and western blot analysis. As shown in Fig. 4, immunofluorescence analysis revealed a significant increase in claudin-1 expression in podocytes of HFD mice compared with ND mice. Consistent with previous reports[16], HFD mice also exhibited a significant elevation of phospho-mTOR in podocytes compared with ND mice. PDS-0330 did not alter claudin-1 expression levels in either ND or HFD mice. However, interestingly PDS-0330 treatment resulted in a significant reduction of phospho-mTOR in podocytes of HFD mice compared with the control group. As shown in Fig. 5, western blot analysis also demonstrated a significant increase in claudin-1 expression in HFD mice compared with ND mice. Consistent with previous reports[16] [9], HFD mice exhibited significant elevations in phospho-Src/total Src, phospho-Akt/total Akt, and phospho-mTOR/total mTOR ratios compared with ND mice. PDS-0330 did not alter claudin-1 expression levels in either ND or HFD mice. Notably, western blot analysis similarly revealed that, in HFD mice, PDS-0330 treatment led to significant reductions in phospho-Src/total Src, phospho-Akt/total Akt, and phospho-mTOR/total mTOR ratios compared with the control group.

**Figure 4.**
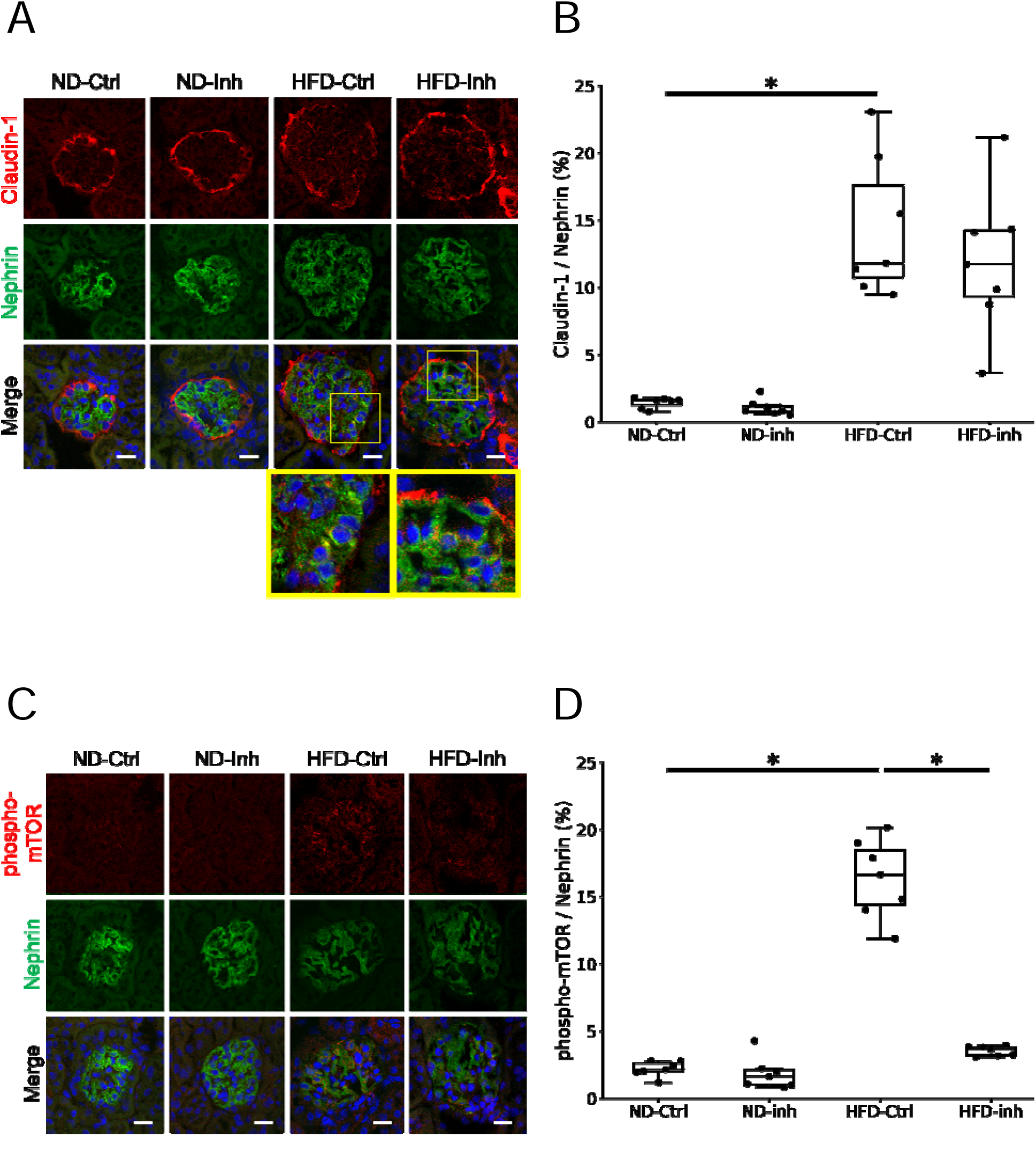
Immunofluorescence staining of glomeruli in mice with or without claudin-1 inhibitor Treatment. (A) Immunofluorescence analysis of claudin-1 expression on podocytes. Podocytes were identified by nephrin staining. Scale bars = 20 μm. (B) Quantification of claudin-1-positive podocyte areas, calculated as the ratio of claudin-1-positive area to the nephrin-positive podocyte area. (C) Immunofluorescence analysis of mTOR phosphorylated at serine 2448 (S2448) on podocytes. Scale bars = 20 μm. (D) Quantification of phospho-mTOR (S2448)-positive podocyte areas, calculated as the ratio of phospho-mTOR (S2448)-positive area to the nephrin-positive podocyte area. Results are presented as the mean ± standard deviation. Differences were evaluated by two-way ANOVA followed by Tukey–Kramer test (* p < 0.05). ND, normal diet; HFD, high-fat diet; Ctrl, control solvent; Inh, claudin-1 inhibitor PDS-0330.

**Figure 5.**
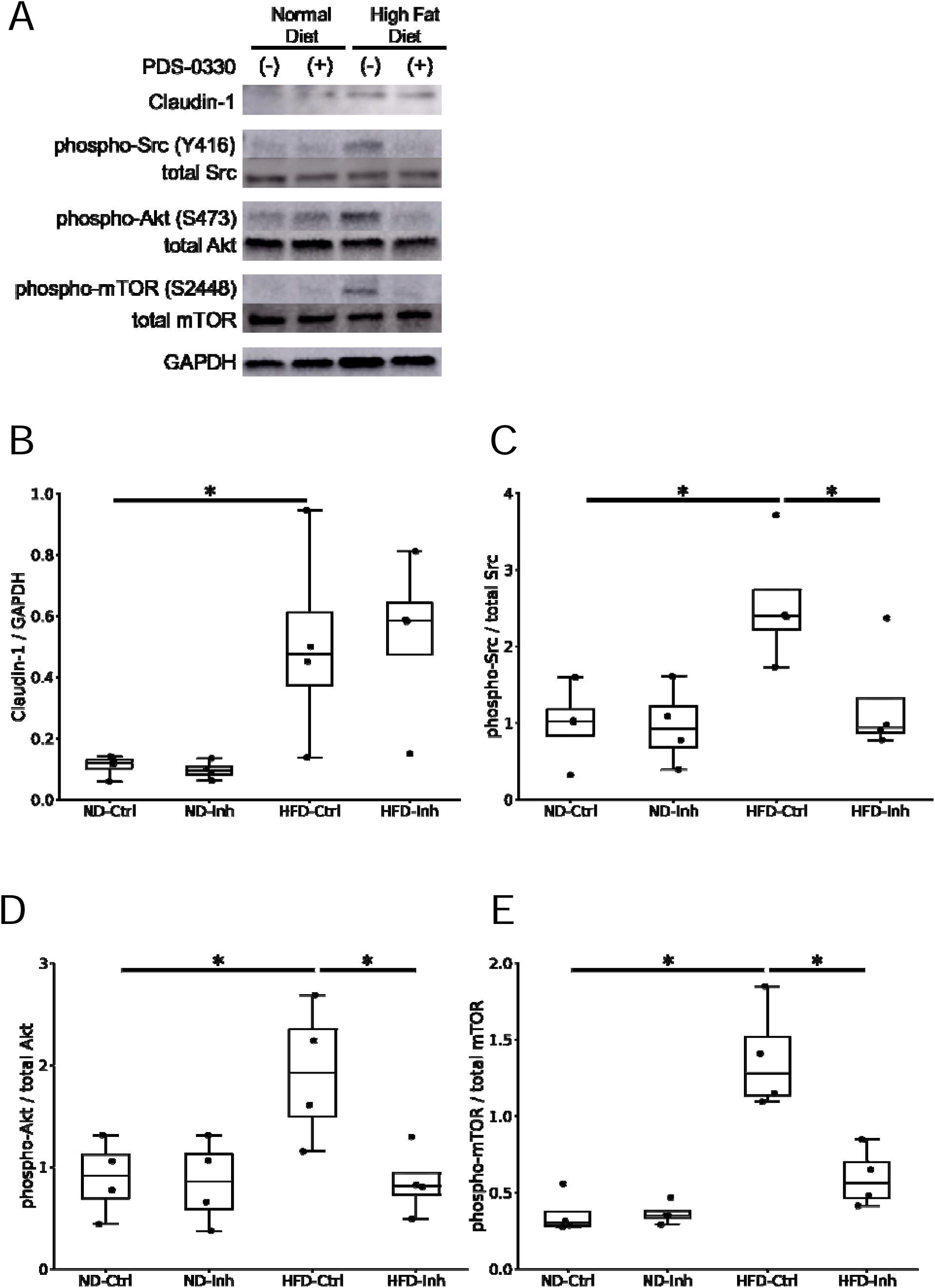
Western blotting analysis of Src, Akt, and mTOR signaling in mouse kidneys treated with or without claudin-1 inhibitor. (A) Western blotting analysis of claudin-1, Src phosphorylated at tyrosine 416 (Y416), Src, Akt phosphorylated at serine 473 (S473), Akt, mTOR phosphorylated at serine 2448 (S2448), mTOR, and Glyceraldehyde-3-phosphate Dehydrogenase (GAPDH). (B-E) Densitometric analysis was performed to quantify the western blotting results. Results are presented as the mean ± standard deviation. Differences were evaluated by two-way ANOVA followed by Tukey–Kramer test (* p < 0.05). ND, normal diet; HFD, high-fat diet; Ctrl, control solvent; Inh, claudin-1 inhibitor PDS-0330.

### PDS-0330 attenuates Src/Akt/mTOR signaling activation in immortalized podocytes under high-glucose conditions

We further evaluated the effects of PDS-0330 in immortalized podocytes using western blot analysis. The immortalized podocyte cell line we used in this study has previously been reported to express claudin-1 in response to adriamycin treatment[17]. As an *in vitro* counterpart of the diabetic model, podocytes were incubated in high glucose (25 mM) for 48 hours, as widely reported in previous studies[18] [19]. To exclude the effects of osmotic stress caused by high-glucose medium, we added an equivalent amount of mannitol to the medium and, consistent with previous reports[20, 21], we confirmed that no significant changes in Src/Akt/mTOR signaling occurred compared with normal glucose conditions (Sup Fig. 1). The treatment concentration of PDS-0330 was set at 25 μM as described in the Materials and Methods section. Then we performed trypan blue staining and confirmed that 25 μM PDS-0330 did not induce any apparent increase in podocyte cell death (Sup Fig. 2). Western blot analyses demonstrated that immortalized podocytes upregulate claudin-1 under high-glucose conditions (Fig. 6). Consistent with the findings in the mouse model, PDS-0330 did not significantly alter claudin-1 expression levels in immortalized podocytes. Interestingly, under high-glucose conditions, PDS-0330 treatment significantly reduced the phospho-Src/total Src, phospho-Akt/total Akt, and phospho-mTOR/total mTOR ratios compared with the control group.

**Figure 6.**
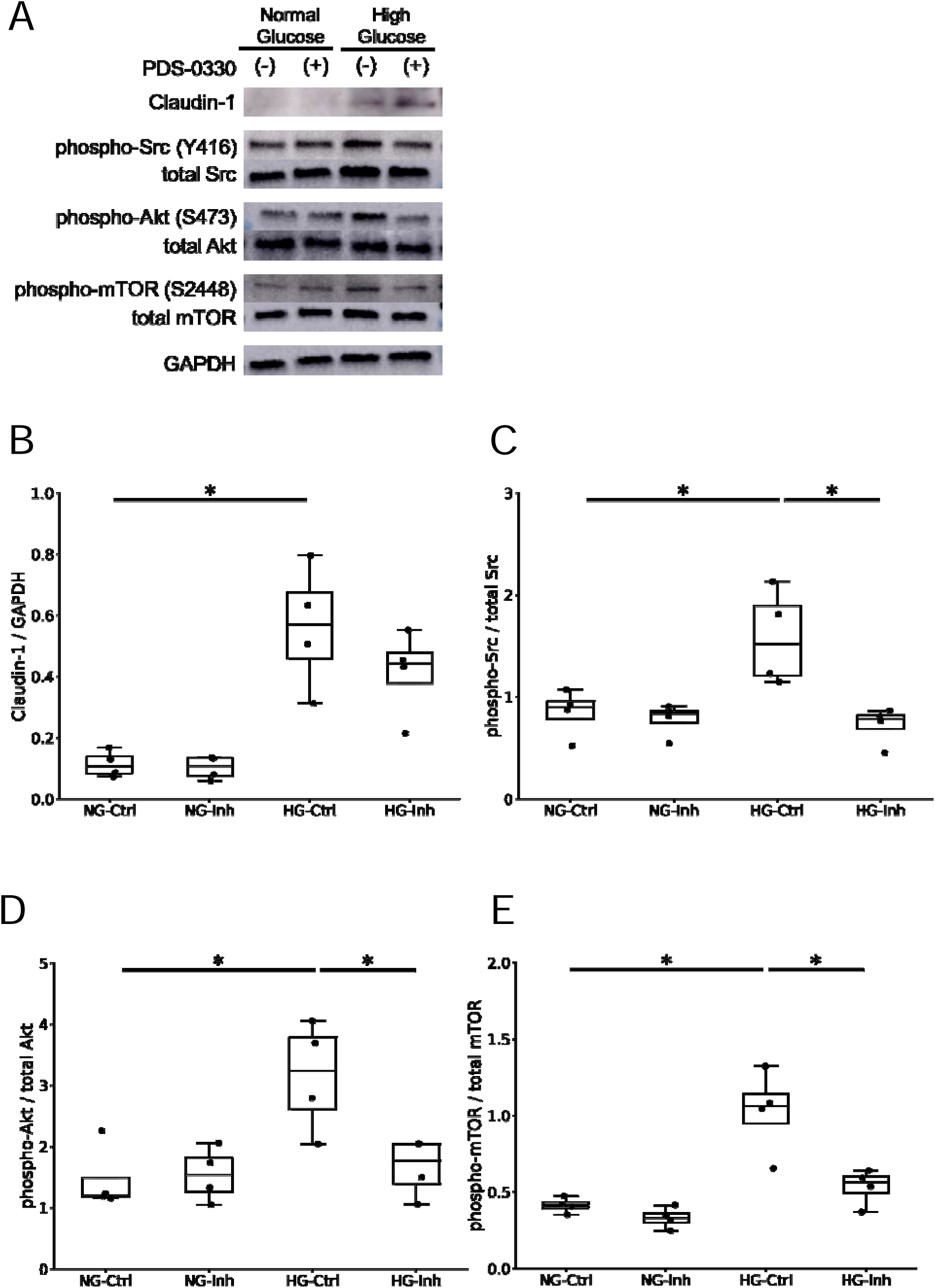
Western blotting analysis of Src, Akt, and mTOR signaling in immortalized podocytes treated with or without claudin-1 inhibitor. (A) Western blotting analysis of claudin-1, Src phosphorylated at tyrosine 416 (Y416), total Src, Akt phosphorylated at serine 473 (S473), total Akt, mTOR phosphorylated at serine 2448 (S2448), total mTOR, and glyceraldehyde-3-phosphate dehydrogenase (GAPDH). (B-E) Densitometric analysis was performed to quantify the western blotting results. Results are presented as the mean ± standard deviation. Differences were evaluated by two-way ANOVA followed by Tukey–Kramer test (* p < 0.05). NG, normal glucose; HG, high glucose; Ctrl, vehicle control; Inh, claudin-1 inhibitor PDS-0330.

Collectively, these results suggest that PDS-0330 exerts therapeutic effects in DKD by suppressing claudin-1/Src/Akt/mTOR signaling activation in podocytes.

## Discussion

In this study, we demonstrated that the claudin-1 inhibitor PDS-0330 exhibits renoprotective effects in a mouse model of DKD. 24-hour urine collection test revealed that PDS-0330 reduced urinary albumin excretion independent of serum glucose levels or body weight, and transmission electron microscopy showed that PDS-0330 exerted protective effects on podocytes. Furthermore, western blot analyses of mouse renal cortical samples revealed that PDS-0330 suppressed Src/Akt/mTOR signaling. The notion that PDS-0330 acts directly on podocytes was further supported by immunofluorescence staining of renal frozen sections and by *in vitro* western blot analyses using immortalized podocyte cell lines. Collectively, these findings suggest that the claudin-1 inhibitor PDS-0330 reduces albuminuria of DKD by acting on podocyte-expressed claudin-1 and inhibiting the Src/Akt/mTOR signaling pathway.

The importance of claudin-1 in the progression of DKD has long been recognized. Previous studies have reported that decreased SIRT1 expression in proximal tubular cells during DKD induces claudin-1 expression in podocytes, leading to albuminuria[6]. It has also been shown that overexpression of SIRT1 reduces podocyte claudin-1 expression and ameliorates albuminuria[7]. These findings indicate that claudin-1 disrupts SIRT1-mediated renoprotective signaling; however, whether claudin-1 itself could serve as a therapeutic target has remained unclear. Our present findings suggest that claudin-1 expressed in podocytes may indeed represent a viable therapeutic target in DKD. A phase II clinical trial of anti-claudin-1 antibody in rapidly progressive glomerulonephritis (RENAL-F02, NCT06047171) is currently underway. In the clinical trial, the anti–claudin-1 antibody is presumed to target cellular crescents within the glomerulus, primarily parietal epithelial cells that constitutively express claudin-1. Whether this antibody agent can also act on podocytes in DKD is an intriguing question. Importantly, the glomerular capillary loop possesses barrier functions that restrict protein passage, and the delivery of immunoglobulin-based therapeutics, including antibody agents, to podocytes may require disruption of this barrier. In contrast, PDS-0330 used in our study is a small-molecule compound with a molecular weight of 423.49 Da, which may allow it to reach podocytes more easily even at early DKD stages characterized only by microalbuminuria. Future studies are needed to determine whether anti–claudin-1 antibody agents and PDS-0330 differ in their efficacy or mechanisms of action.

In the field of oncology, claudin-1 has been shown to influence multiple signaling pathways and is recognized as one of the master regulators of cancer cell invasion and proliferation[3]. Among the downstream target proteins of claudin-1 signaling is Src, which plays a critical role not only in cancer progression but also in the development of DKD[9]. Furthermore, mTOR, which is positioned downstream of Src, is also a key molecule in the progression of DKD. Indeed, both Src inhibitors and mTOR inhibitors have been reported to attenuate the progression of DKD[9] [16], and our present findings are consistent with these reports. These inhibitors, however, are associated with various systemic adverse effects[22] [23] [24] [25], partly because Src and mTOR are expressed broadly and non-specifically across many cell types. In contrast, PDS-0330 used in our study is designed to act specifically on cells that reactively express non-junctional claudin-1, suggesting the possibility of fewer off-target effects. Given that non-junctional claudin-1 affects diverse signaling pathways and is selectively expressed in pathological cells, the development of DKD therapies targeting non-junctional claudin-1 may represent a promising therapeutic strategy. Of course, it should be noted that PDS-0330, like other anticancer agents, might also induce cell death and cause adverse effects. The factors determining cellular sensitivity to PDS-0330 remain to be elucidated. The absence of PDS-0330–induced cell death in podocytes, despite claudin-1 expression, may reflect fundamental differences in claudin-1 dependency and expression levels between podocytes and cancer cells. Cancer cells often depend on aberrant Src signaling for survival[26], whereas podocytes rely on multiple parallel survival pathways and express claudin-1 at much lower levels. These differences might explain why PDS-0330 induces anoikis, a form of programmed cell death, in cancer cells but not in podocytes. Further studies are needed to identify the factors that determine whether claudin-1–targeted therapy induces cell death before such treatments can be applied in clinical practice.

Our study has at least one limitation. We didn’t completely exclude the possibility that PDS-0330 acts on targets other than claudin-1 on podocyte. However, our experimental findings support the specificity of PDS-0330 for claudin-1: podocytes cultured under normal-glucose conditions, in which claudin-1 expression is very few, showed almost no change in Src activation upon PDS-0330 treatment, whereas podocytes cultured under high-glucose conditions — with markedly increased claudin-1 expression — exhibited significant Src suppression (Fig. 6). This strongly suggests that the claudin-1– dependent inhibition of Src activity — an intrinsic property of PDS-0330

— was indeed exerted via podocyte-expressed claudin-1.

In this study, we didn’t elucidate the mechanism by which PDS-0330 reduced albuminuria even in ND mice. However, it is worth noting that PDS-0330 may have exerted a direct protective effect on podocytes in ND mice as well. This possibility is supported by our observations that ND mice, despite having urinary albumin levels consistent with previous reports[10] [11], still exhibited low-level claudin-1 expression in podocytes and showed segmental foot process effacement. It is well known that healthy mice exhibit higher levels of urinary albumin than healthy humans, largely because mice possess far fewer glomeruli than humans and therefore experience a greater workload per glomerulus[10]. In addition, testosterone has been reported to increase albuminuria in male mice[10]. These factors may impose stress on podocytes, potentially inducing claudin-1 expression and contributing to albuminuria even under ND conditions. Such findings raise the possibility that claudin-1–targeted therapies may be effective not only in DKD but also in podocyte injuries arising from other mechanisms. In this study, PDS-0330 did not significantly alter Src/Akt/mTOR signaling in ND mice. This might be because the baseline activity of these signaling pathways is inherently low in normal mice, making subtle changes difficult to detect by western blotting, or this might be because PDS-0330 may have influenced other signaling pathways instead. Further studies are needed to determine which additional signaling pathways are modulated by claudin-1–targeted therapy.

In conclusion, the claudin-1 inhibitor PDS-0330 exerts renoprotective effects in DKD by acting on podocytes and suppressing Src/Akt/mTOR signaling. Further studies of claudin-1–mediated pathogenic mechanisms, as well as the development of therapeutic strategies targeting claudin-1, will contribute to the advancement of DKD treatment.

## Material and Methods

### Animal Models

This study was approved by the animal care and use committee of Okayama University (OKU-2023833). We used five-week-old male mice with a C57BL/6J genetic background (CLEA Japan, Tokyo, Japan). The mice were fed with normal diet with the composition of 13 kcal% fat (Cat#: MF, Oriental Yeast Co., ltd, Tokyo, Japan) or high-fat diet with the composition of 60 kcal% fat (Cat#: D12492, Research Diets INC, New Brunswick, USA). After ten weeks of feeding, we orally administrated 5 mg/kg PDS-0330 (MedChemExpress LLC, Monmouth Junction, NJ, USA) dissolved in a solvent consisting of 5% dimethyl sulfoxide (DMSO), 40% polyethylene glycol 300 (PEG300), 5% polyoxyethylene (20) sorbitan monooleate (Tween-80) and 50% double-distilled water or the solvent (as control) once weekly for 4 weeks. After that, the mice were sacrificed, and the serum plasma, urine (collected for 24 hours), and kidneys were obtained.

### Cell Culture

As previously described[27], conditionally immortalized mouse podocytes, provided from Dr. Peter Mundel[28], were cultured on collagen type I (BD Biosciences, San Jose, CA, USA)-coated dishes and maintained at the permissive condition of 33 °C in Roswell Park Memorial Institute (RPMI) 1640 medium (Thermo Fisher Scientific, Waltham, MA, USA), supplemented with 10% fetal bovine serum (FBS) (Thermo Fisher Scientific, Waltham, MA, USA), 1% penicillin and streptomycin (Thermo Fisher Scientific, Waltham, MA, USA), and 200 U/mL γ-interferon (Sigma-Aldrich, St. Louis, MO, USA). The podocytes were transferred to a non-permissive condition of 37 °C without γ-interferon for cell differentiation for about 7 days and were used for the further experiment. For cell treatment, podocytes were seeded on 6-well pre-coated plates. Cell starvation was conducted for 24 hours (starvation medium supplemented with 0.5% FBS, 1% penicillin and streptomycin). Then cells were cultured in RPMI 1640 medium, either as supplied or supplemented with 19.5 mM glucose to a final concentration of 25 mM, or 19.5 mM mannitol, and treated with 25 µM PDS-0330 (DMSO vehicle; final DMSO concentration 0.35%) or DMSO as a control for 48 hours. The western blotting was then conducted. The treatment concentration of PDS-0330 was determined based on previously reported findings as below[8]; In prior studies, claudin-1-negative cancer cells did not exhibit a significant increase in cell death at PDS-0330 concentrations up to 20 μM, whereas a significant increase in cell death was observed at 40 μM. In contrast, claudin-1-high-expressing cancer cells showed a significant reduction in Src activation at PDS-0330 concentrations of 12.5 μM and 25 μM. Considering that podocytes might be less sensitive to PDS-0330 than claudin-1-high-expressing cancer cells, and that higher concentrations of PDS-0330 might induce cytotoxicity, we selected 25 μM as the treatment concentration for our experiments.

### Histopathological Analysis

Sections were cut to 4 μm thickness from 10% formalin-fixed paraffin-embedded kidney samples. These sections were used for periodic acid–Schiff staining.

For immunofluorescence microscopy, frozen sections (4 μm) were stained using anti-claudin-1 antibody (1:100, #: 13255S, Cell Signaling Technology, Danvers, MA, USA), anti-nephrin antibody (1:200, #: GP-N2, Progen Biotechnik GmbH, Heidelberg, Germany), anti-phospho mTOR (Ser2448) antibody (1:100, #: 2971, Cell Signaling Technology, Danvers, MA, USA) and 4′,6-diamidino-2-phenylindole (DAPI). Sections were examined using an Olympus FSX100 biological microscope (Olympus Corporation, Tokyo, Japan) or Zeiss LSM780 confocal microscope (Carl Zeiss Microscopy GmbH, Jena, Germany) at the Central Research Laboratory of Okayama University Medical School. In a kidney histological quantitation, a minimum of six non-overlapping images of cortex per kidney were observed under 400× magnification. Quantitative computer-assisted image analysis was performed using a digital image-analyzing software, ImageJ (available at http://rsbweb.nih.gov/ij/index.html; National Institutes of Health).

For electron microscopy, kidney specimens were fixed with 2.5% glutaraldehyde and observed using a transmission electron microscope (H-7650, Hitachi, Tokyo, Japan).

For quantitative analysis of podocyte foot process width (FPW), we adopted a previously described method [29]. Kidneys from 12 mice (4 groups, 7 mice per group; 3 mice per group were randomly selected for electron microscopic analysis) were examined. For each mouse, 4 glomeruli were randomly selected, and 5 open capillary loops were analyzed per glomerulus (total 20 loops per mouse). Electron micrographs were acquired at a nominal magnification of ×5000, and FPW was calculated for each mouse.

### Western Blot Analysis

Proteins were extracted using a lysis buffer (Cat #: 89900, Thermo Fisher Scientific, Inc., Waltham, MA, USA) containing a protease inhibitor (Cat #: G652A, Promega Corporation, Fitchburg, WI, USA) and a phosphatase inhibitor (Cat #: ab201113, Abcam plc., Cambridge, UK), then quantified using a bicinchoninic acid (BCA) protein assay kit (Thermo Fisher Scientific Inc., Waltham, MA, USA). Equal amounts of protein were separated using 10% sodium dodecyl sulfate–polyacrylamide gel electrophoresis and electrophoretically transferred onto nitrocellulose membranes, which were blocked with 1% bovine serum albumin (Cat #: A7888-50G, Sigma-Aldrich Co. LLC, St. Louis, MO, USA) for 30 min. The membranes were incubated with the following primary antibodies overnight at 4 °C: anti-claudin-1 antibody (1:1,000, #: 13050-1-AP, Proteintech Group, Inc., Rosemont, IL, USA), anti-phospho Src (Tyr416) antibody (1:1,000, #: 6943, Cell Signaling Technology, Danvers, MA, USA), anti-Src antibody (1:1,000, #: 2109, Cell Signaling Technology, Danvers, MA, USA), anti-phospho Akt (Ser473) antibody (1:1,000, #: 4060, Cell Signaling Technology, Danvers, MA, USA), anti-Akt antibody (1:1,000, #: 4691, Cell Signaling Technology, Danvers, MA, USA), anti-phospho mTOR (1:1,000, #: 2971, Cell Signaling Technology, Danvers, MA, USA), anti-mTOR antibody (1:1,000, #: 2972, Cell Signaling Technology, Danvers, MA, USA), and anti-glyceraldehyde 3-phosphate dehydrogenase antibody (GAPDH; 1:10,000, #: 2118, Cell Signaling Technology, Danvers, MA, USA). They were incubated with a horseradish peroxidase (HRP)-conjugated secondary antibody (1:2,000, Cat#: 170-6515, Bio-Rad Laboratories, Inc., Hercules, CA, USA) for 1 hour. The membranes were extensively washed in phosphate-buffered saline with Tween-20, and antigen–antibody complexes were visualized by chemiluminescence using an enhanced chemiluminescence (ECL) kit (ECLTM Prime Western Blotting Reagents, GE Healthcare, Chicago, IL, USA).

### Statistical Analysis

Results are presented as the mean ± standard deviation. Differences among groups were evaluated by two-way ANOVA with diet (normal diet vs. high-fat diet) and treatment (control vs. PDS-0330) as fixed factors, followed by Tukey–Kramer multiple comparison test. Statistical analyses were performed using Python (Anaconda Distribution, version 3.x; Anaconda Inc., Austin, TX, USA). Statistical significance was accepted at p < 0.05.

## Author contributions

K.F. conceived and designed research; K.F., K.T., H.N., and N.U. performed experiments; K.F. and K.T. analyzed data; K.F. and K.T. interpreted results of experiments; K.F. and K.T. prepared figures; K.F. drafted manuscript; K.F., K.T., H.N., N.U., S.K., and J.W. edited and revised manuscript; K.F., K.T., H.N., N.U., S.K., and J.W. approved final version of manuscript.

## Acknowledgements

We thank the Central Research Laboratory, Okayama University Medical School, for the assistance with the confocal laser scanning microscope and kidney tissue processing.

