## Supplementary Figures for "Claudin-1 Inhibitor PDS-0330 Ameliorates Diabetic Kidney Disease by Suppressing Src/Akt/mTOR Signaling Pathway in Podocytes"

### Slide 1
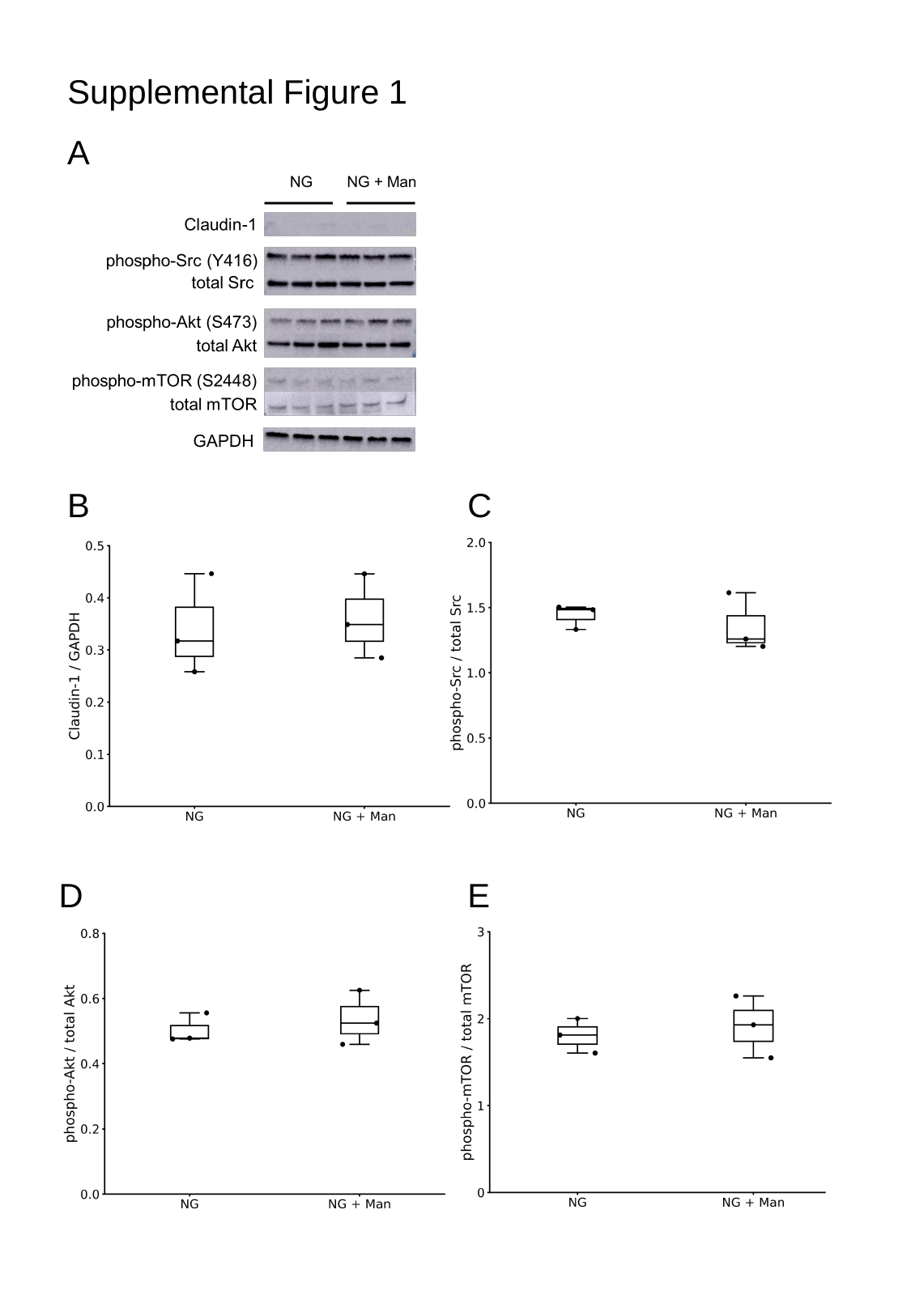

Supplemental Figure 1
A
B
C
D
E

### Slide 2
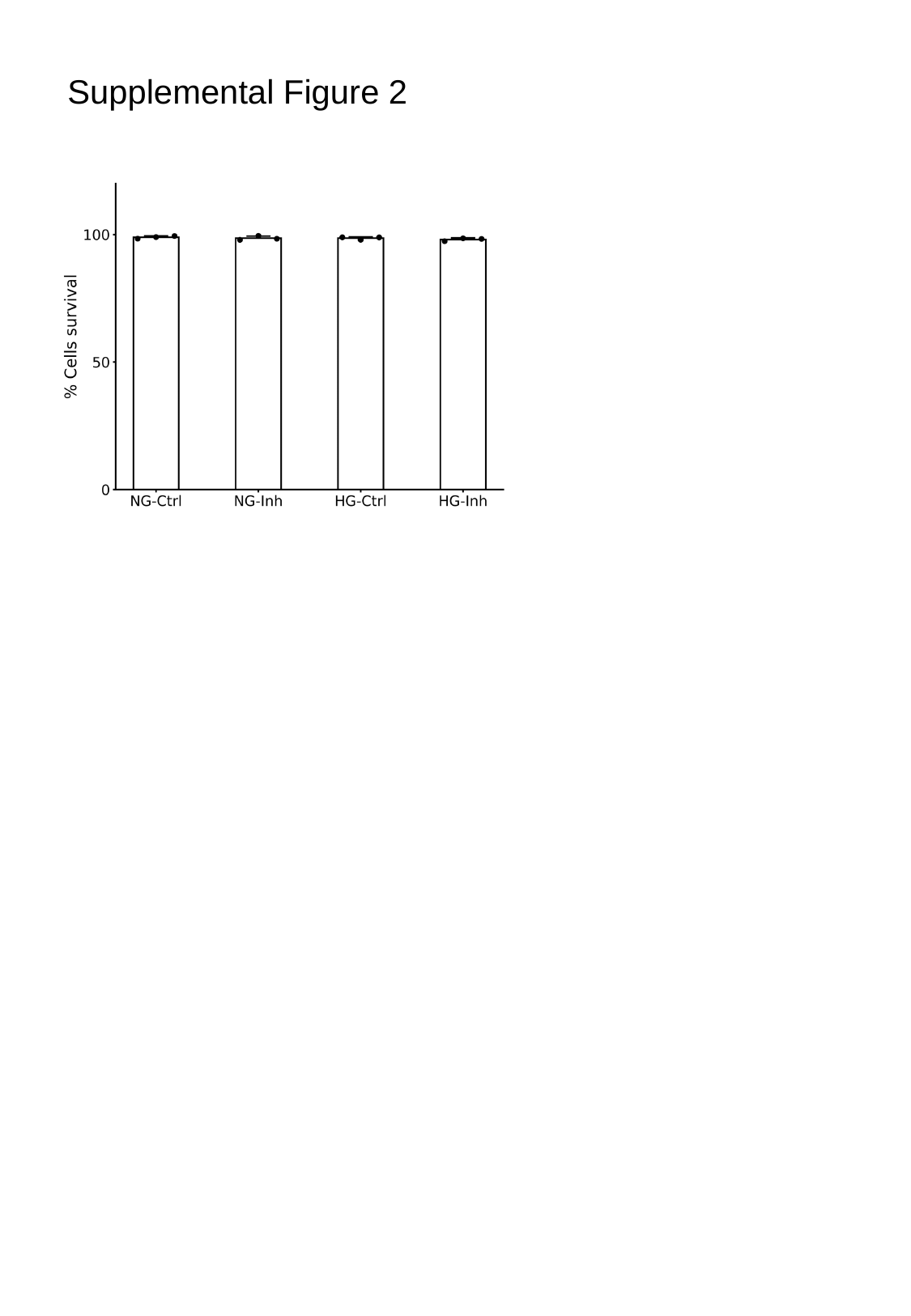

Supplemental Figure 2

### Slide 3
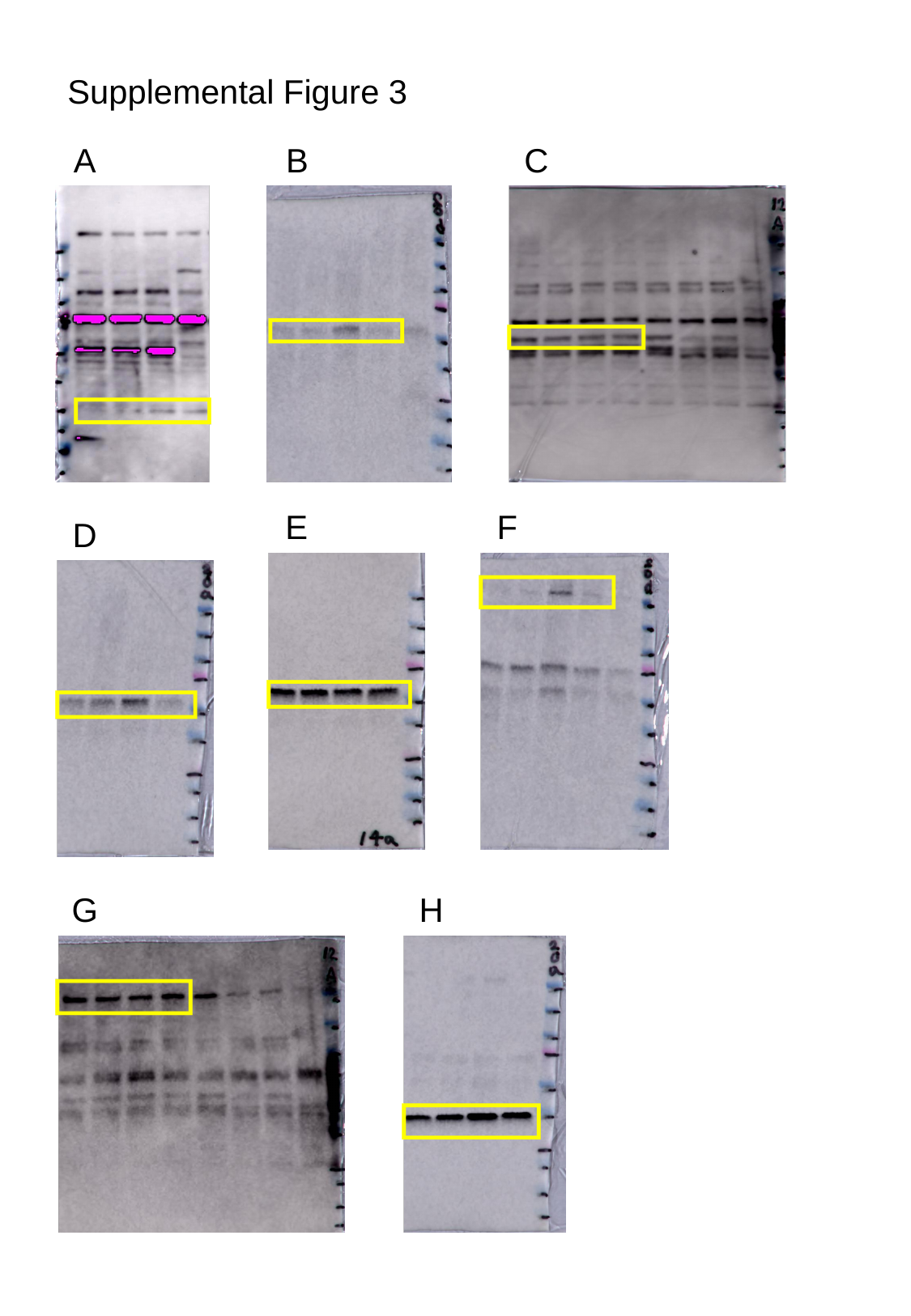

Supplemental Figure 3
A
B
C
E
F
D
G
H

### Slide 4
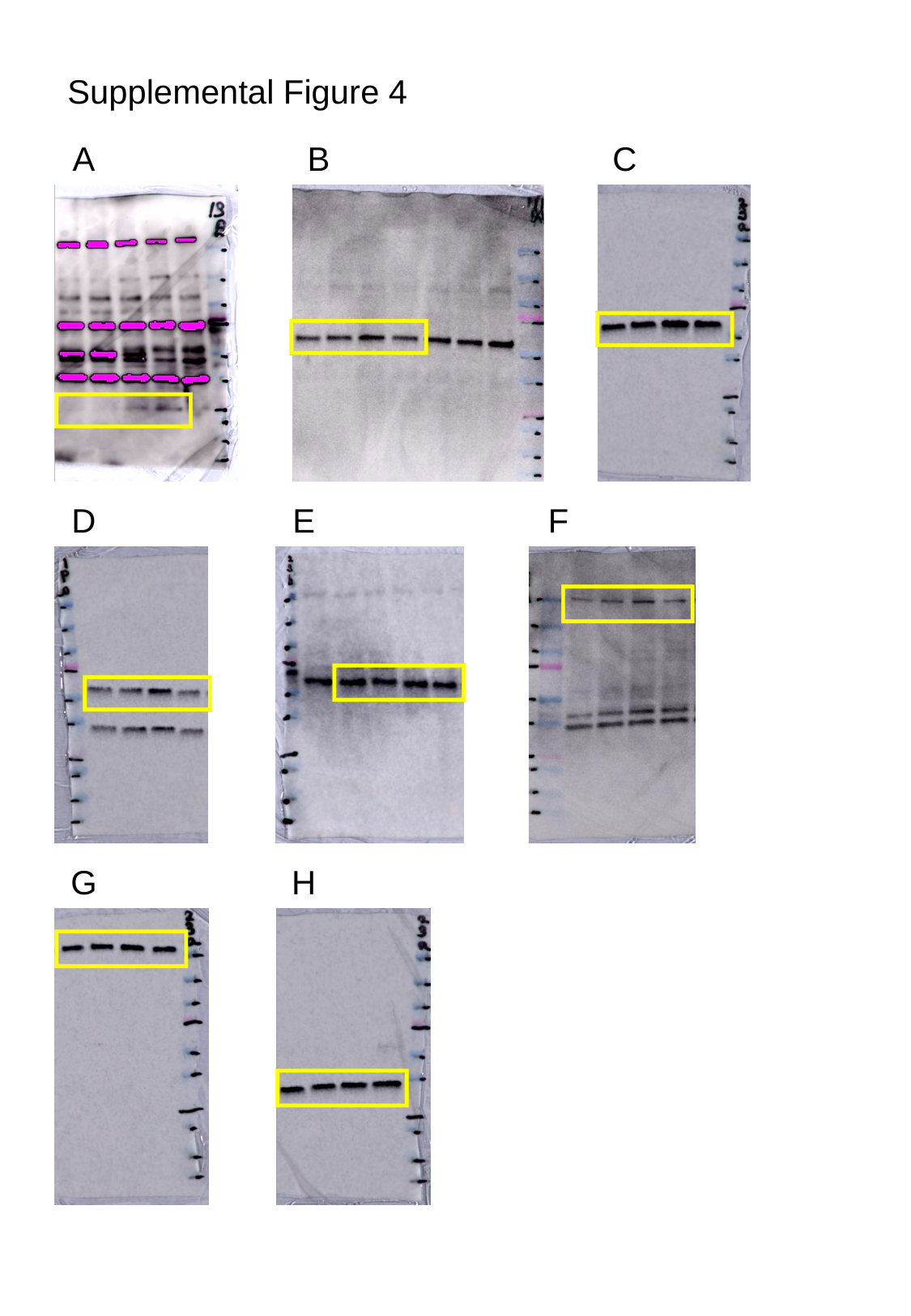

Supplemental Figure 4
A
B
C
D
E
F
G
H

### Slide 5
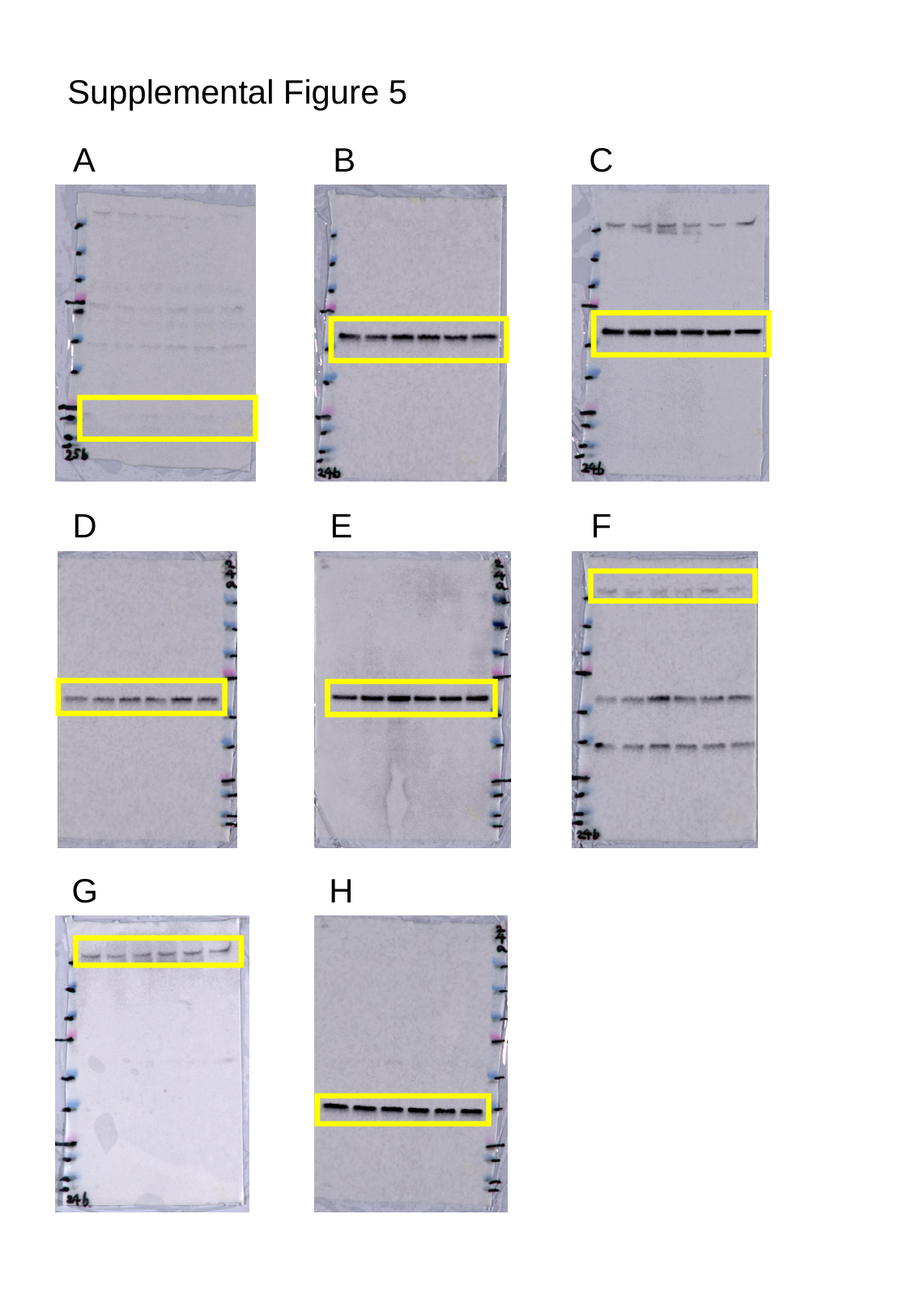

Supplemental Figure 5
A
B
C
D
E
F
G
H
